# L-lactate is a regulator of gametocytogenesis in *Plasmodium falciparum*

**DOI:** 10.64898/2026.09.25.754385

**Authors:** Nana Efua Andoh, Ibtissam Jabre, Catherine J. Merrick

**Affiliations:** Department of Pathology, University of Cambridge, Tennis Court Road, Cambridge, CB2 1QP, UK

**Keywords:** Malaria, *Plasmodium*, gametocytogenesis, lactate, histone lactylation

## Abstract

Epigenetic processes play important roles in the biology of the malaria parasite *Plasmodium falciparum*. We recently showed that *Plasmodium* histones are not only acetylated and methylated but also lactylated. This new epigenetic mark could form a particularly important axis of host-parasite signalling because severe malaria is often characterised by hyperlactataemia. In the human host this can cause respiratory distress, which is potentially fatal. For the parasite, it could be advantageous to sense this state of pathology and respond by modulating virulence. Virulence processes known to be under epigenetic control include antigenic variation, invasion switching and conversion to sexual gametocytes, i.e. gametocytogenesis. Here, we investigated the influence of lactate on gametocytogenesis. Hyperlactataemia in malaria patients is defined as ≥ 5mM blood lactate and we confirmed that in laboratory culture, adding 5mM L-lactate was sufficient to boost gametocytogenesis. Expression of the *GDV-1* gene increased within one cell cycle of asexual parasites being exposed to L-lactate, suggesting that the canonical epigenetic switch for gametocytogenesis was involved: GDV-1 regulates epigenetic de-repression of the gene encoding the master transcription factor for gametocytogenesis, AP2-G. Indeed, parasite histones became rapidly lactylated after L-lactate exposure and chromatin profiling by CUT&Tag detected inducible lactylation of chromatin specifically upstream of the *AP2-G* gene, and also of the antisense RNA that regulates *GDV-1*. Thus L-lactate joins S-adenosylmethionine as a metabolite that can epigenetically regulate gametocytogenesis in *P. falciparum*.

**Author summary:** Malaria in humans often causes elevated blood lactate: a phenomenon called hyperlactataemia. This occurs because, during malarial disease, both malaria parasites and human tissues tend to respire by glycolysis instead of oxidative respiration. For bloodstream parasites this is their only mode of respiration, while human tissues become hypoxic when parasitized cells adhere in capillaries and impede the flow of oxygenated blood. Hyperlactataemia predicts severe and fatal malarial disease because it causes respiratory distress.

Here, we show that malaria parasites in culture can sense and respond to elevated lactate – mimicking hyperlactataemia – by increasing their conversion to sexual cells, which are vital for transmission to mosquitoes. When a human host is at risk of death, it could be advantageous for the parasite to boost its chances of transmission thus. The switch occurs by increasing expression of genes that promote sexual conversion: genes that are known to be silenced or activated epigenetically. Epigenetic changes are chromatin marks that are made without changing the underlying DNA sequence: they are fast, flexible and reversible, making them ideal for responding to changing situations in a human host. Thus, a blood metabolite that is characteristically raised during malaria can epigenetically influence an important virulence phenotype in malaria parasites.

## Introduction

*Plasmodium falciparum* is the most important cause of human malaria, responsible for over half a million malaria deaths each year [1]. This parasite has many unusual features in its biology and life cycle, including the way it transmits to its mosquito vector. Bloodstream parasites usually convert a proportion of their mitotically-dividing cells into presexual cells called gametocytes, which exit the mitotic cycle, differentiate irreversibly, and circulate in a quiescent state until they are taken up by a mosquito bite. They then develop into mature gametes and mate in the mosquito gut.

In most *Plasmodium* species, developmental conversion to gametocytes takes just a few days and the resultant cells are not easily distinguishable from asexual cells. In *P. falciparum*, conversion takes ∼2 weeks, progresses through several morphologically distinct stages and culminates in distinctive crescent-shaped gametocytes. This protracted development occurs not in circulating blood but sequestered primarily in bone marrow [2, 3].

At the molecular level, gametocytogenesis requires an epigenetic switch, turning on the expression of the master transcription factor AP2-G [4, 5]. This is regulated in turn by blood levels of lysophosphatidylcholine, which influence the level of S-adenosylmethionine in the parasite and hence the efficiency of histone methylation, which epigenetically silences *AP2-G* [6]. Other factors, however, may also influence gametocytogenesis [7] and most of them are biochemically and mechanistically ill-defined. For example, elevated lactic acid has been reported to boost gametocytogenesis in culture: adding 8.2mM lactic acid to the daily media supplied to a culture during developmental conversion resulted in 1.5-fold higher gametocytaemia after 2 weeks [8].

We recently reported that exposing *P. falciparum* to elevated lactate in culture stimulated the appearance of a novel epigenetic mark, histone lactylation [9, 10]. Therefore, the genes that control gametocytogenesis could theoretically be regulated not only by methylation, influenced by the metabolite lysophosphatidylcholine, but also by lactylation, influenced by the metabolite lactate. The previous publication on this topic [8] did not define a molecular mechanism for the effect of lactic acid, and it raised other questions as well. Firstly, the lactic acid used was a mixture of L- and D-forms, reportedly containing only ∼3.1mM L-lactate [8], whereas the biologically-relevant form is overwhelmingly L-lactate [11, 12]. Secondly, only a single level of lactic acid exposure was tested, which may or may not be optimal: the clinical threshold for hyperlactataemia is 5mM, while the blood of severe malaria patients can reach at least 15mM [13]. Thirdly, it was unclear whether lactic acid exclusively stimulated the developmental commitment of asexual cells, or whether it also enhanced gametocyte maturation. Fourthly and most importantly, no molecular mechanism was reported: *AP2-G* expression was not tested, and other genes expressed in early gametocytes were reportedly unchanged.

Here, we set out to address these questions, and to establish whether the epigenetic pathway of histone lactylation influences *P. falciparum* gametocytogenesis.

## Results

### Moderate levels of L-lactate induce gametocytogenesis in cultured *P. falciparum*

We reproduced the published result [8] using defined concentrations of sodium L-lactate. Parasites were exposed to 5mM added L-lactate throughout a 2-week gametocytogenesis protocol (Fig 1A), resulting in elevated gametocytaemia (Fig 1B). This was modest, ∼1.5-fold, but the elevation was consistent across 4 biological replicates and comparable to the 1.5-fold increase caused by 8.2mM lactic acid (∼3.1mM L-lactate) [8]. There was a trend towards improved results with 10mM L-lactate (Fig 1C) and we therefore tested a variant protocol, starting with 4% rather than 0.5% asexual parasitaemia, because higher parasitaemias would add more endogenously-generated lactate to the culture, potentially boosting the effect. In fact, however, added L-lactate had no detectable effect in this protocol (Fig 1D). We observed that at lower parasitaemia (as used in [8]), asexual cells survived for several cycles before growing to a lethal hyperparasitaemia that left only maturing gametocytes in the culture. Higher parasitaemias, by contrast, precipitated a faster ‘crash’, hence the time available for abundant asexual cells to commit was presumably reduced.

**Fig 1:**
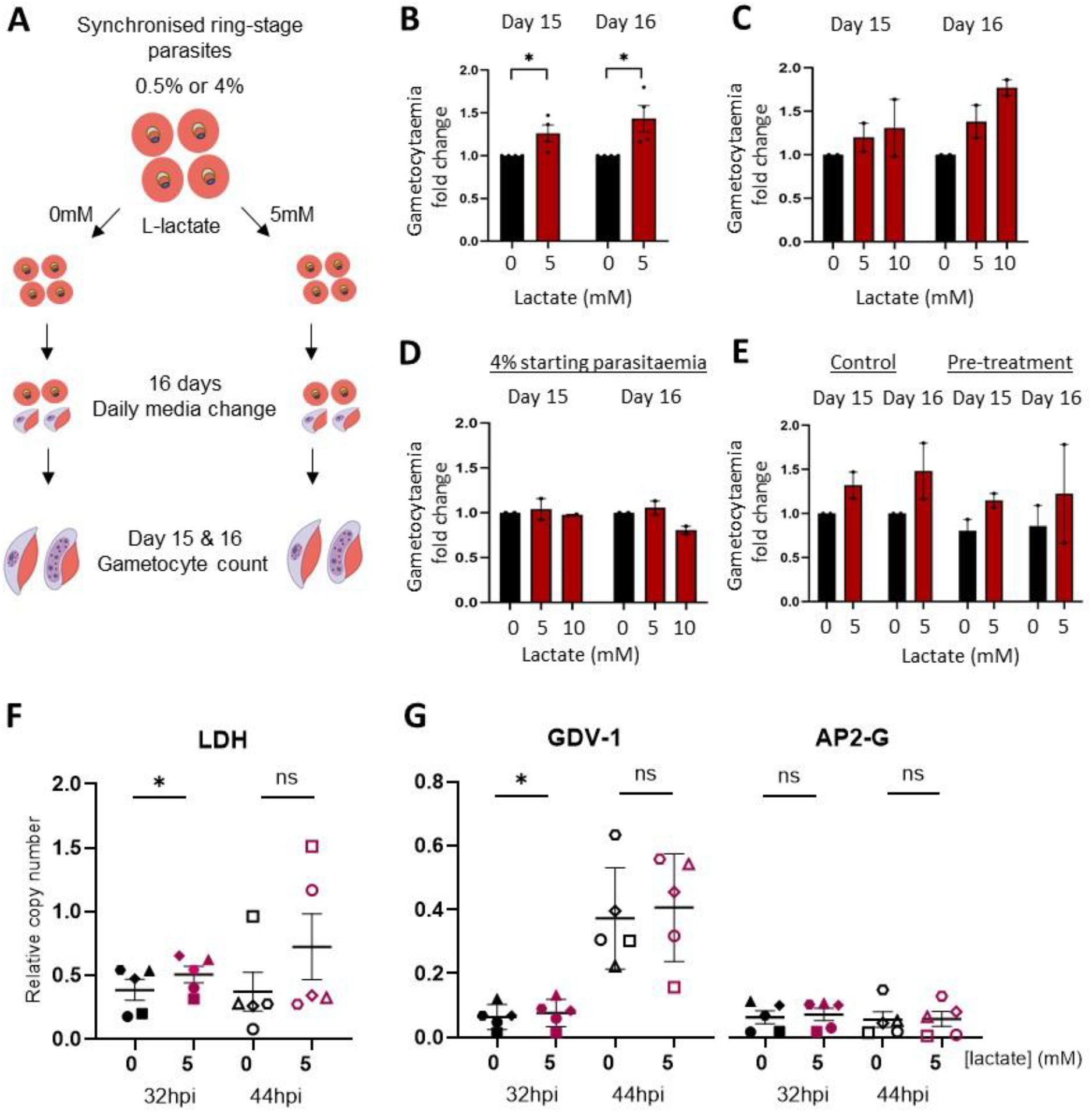
Moderate levels of L-lactate induce gametocytogenesis and *GDV-1* gene expression in cultured *P*. *falciparum*. A) Schematic showing the protocol for gametocyte induction with or without added L-lactate. B) Mature stage-V gametocytes were counted at days 15 and 16 in 4 independent biological replicate experiments conducted +/-5mM L-lactate. Gametocytaemia in each lactate-treated condition was expressed as a fold-change from the gametocytaemia in the corresponding control culture, then plotted as mean +/-SEM. Significance was tested via paired one-tailed T-test: *, p < 0.05. C) Mature stage-V gametocytes were counted at days 15 and 16 in 2 independent biological replicate experiments with 0, 5 or 10mM L-lactate. Data plotted as in (B); significance tested via repeated measures one-way ANOVA: no comparisons reached significance at p < 0.05. D) Mature stage-V gametocytes were counted at days 15 and 16 in 2 independent biological replicate experiments conducted from 4% parasitaemia (rather than 0.5%, as in B, C) with 0, 5 or 10mM L-lactate. Data plotted as in (B); significance tested via repeated measures one-way ANOVA: no comparisons reached significance at p < 0.05. E) Asexual cultures at 0.5% parasitaemia were first ‘pre-treated’ +/-5mM L-lactate for 2 growth cycles, before commencing the protocol shown in (A), treating +/-5mM L-lactate throughout 16 days. Mature stage-V gametocytes were counted at days 15 and 16 in 2 independent biological replicate experiments. Data plotted as in (B), expressing the gametocytaemia in all conditions in each experiment as a fold-change from the gametocytaemia in the corresponding control culture. Significance tested via repeated measures two-way ANOVA: no comparisons reached significance at p < 0.05. F) Expression levels of *LDH* were measured by RT-qPCR at 32 and 44h after treatment of synchronised asexual cultures +/-5mM L-lactate. Relative copy number of *LDH* was calculated versus the mean expression level of 4 housekeeping genes. Values from 5 independent biological replicate experiments are shown, with mean and SD. Significance was tested via paired one-tailed T-test: *, p < 0.05; ns, non-significant. G) Expression levels of *GDV-1* and *AP2-G* were measured and plotted as in (F). The mean relative copy number of *GDV-1* after 0 vs. 5mM L-lactate was 0.064 vs. 0.076 at 32h; 0.37 vs. 0.41 at 44h. Mean relative copy numbers of *AP2-G* were 0.063 vs. 0.072 at 32h; 0.056 vs. 0.057 at 44h.

### L-lactate exposure throughout the period of gametocytogenesis is the most effective regime to boost gametocytaemia

We next sought to distinguish the influences of L-lactate on gametocyte commitment versus maturation. Prior results [8] suggested that both could be affected, because adding lactic acid for only the first 8 days of a 16-day protocol increased gametocytaemia less than adding it for the full 16 days. Over only the final 8 days, lactic acid boosted gametocyte numbers even less (presumably because fewer surviving asexual cells were available to commit by day-9), but it did slightly increase the gametocytes’ mosquito infectivity.

Since days 1-8 encompass both the commitment of asexual cells and the maturation of early-stage gametocytes, we designed an experiment to better isolate commitment. We exposed parasites to 5 or 10mM L-lactate *only* during 2 cycles of asexual growth from a low 0.5% parasitaemia, before starting the 16-day protocol of progressive over-growth with or without added lactate (as per Fig 1A). We hypothesised that ‘pre-treatment’ alone would boost gametocyte numbers if it was sufficient to induce commitment. In fact, ‘pre-treatment’ with 5mM L-lactate had no detectable effect (Fig 1E), and ‘pre-treatment’ with 10mM L-lactate prevented asexual parasite growth when it was maintained over more than one 48h cycle, thus precluding a downstream gametocytogenesis assay. We concluded that L-lactate exposure *within* the overgrowth protocol was required to achieve a ∼1.5-fold increase in gametocytaemia.

### L-lactate exposure swiftly increases expression of the *GDV-1* gene that regulates gametocytogenesis

The master regulator of gametocyte commitment, AP2-G [4, 5], is itself regulated by GDV-1 [14], expressed within the asexual growth cycle when commitment occurs [15] (well before gametocyte morphology appears). To determine whether this genetic pathway was promptly induced through L-lactate, we harvested RNA during a single cycle of exposure to 5mM lactate and measured gene expression at two timepoints: in trophozoites at 32 hours post invasion (hpi) and in schizonts at 44 hpi. As a control, we also measured lactate dehydrogenase (*LDH*) expression, because elevated lactate levels were likely to induce this gene.

*LDH* was indeed induced, ∼1.3-fold in trophozoites and almost 2-fold in schizonts, albeit with considerable variation between replicates (Fig 1F). *GDV-1* was also induced (Fig 1G), reaching ∼1.2-fold with statistical significance in trophozoites, but not in schizonts, which again showed more inter-replicate variation. Within this acute timeframe, *AP2-G* transcription remained very low and was not significantly induced (Fig 1G). Thus, *GDV-1* can apparently undergo an induction cascade in response to lactate within one cell cycle, whereas it may take longer for significant changes in *AP2-G*. Low overall expression levels of these genes were expected because only a minority of cells in any culture were likely to be committing. High variation between five independent experiments was also expected because other uncontrollable factors, such as blood and serum batches, strongly influence the overall rate of commitment. Nevertheless, since we detected acute and consistent *GDV-1* induction, the lactate signal is probably transduced through the classical *GDV-1*/*AP2-G* expression pathway [14].

### L-lactate exposure swiftly increases histone lactylation upstream of the *AP2-G* gene

We then sought evidence that gene induction was epigenetic, i.e. that it occurred through lactylation of histones. We hypothesised that adding L-lactate should increase the parasite’s pool of available lactyl moieties, and hence the level of histone lactylation, just as S-adenosylmethionine increases the pool of methyl donors for histone methylation [6]. Lactylation is generally considered gene-activating, similarly to acetylation [16], although this code is not yet well characterised.

We previously showed that adding lactate at 5mM or more to an asexual culture for 12h could increase histone lactylation in parasite chromatin [10]. The effect was robust and titratable: pulses of up to 25mM lactate caused progressively stronger histone lactylation [10]. In Figure 2A, we show that lactylation is not only titratable but highly temporally dynamic, being boosted in trophozoite chromatin within just 2h of a strong 25mM-lactate stimulus, sustained for at least 12h of lactate exposure, and reversible within 6h of washout (Fig 2A). Based on these parameters, we used Cleavage Under Targets and Tagmentation (CUT&Tag) to focus upon histone lactylation specifically within the *AP2-G* and *GDV-1* genes. Both genes showed lactylation, primarily within the coding sequence, consistent with the pattern we previously observed genome-wide [17] (Fig 2B, C). The upstream region of *AP2-G*, which was otherwise minimally lactylated, became inducibly marked after 12h of trophozoite exposure to 25mM lactate: this occurred on at least 2 histones, H3K18 and H4K12, whereas H4 acetylation was not induced (Fig 2B). Again, only a small minority of cells in the population were likely to be committing, so the signal may be even stronger within committing cells. *GDV-1* by contrast was not similarly marked (Fig 2C), but a small peak of KLa did appear at the end of the antisense RNA that controls *GDV-1* expression, and a small peak of H4 acetylation appeared upstream of the gene itself.

**Fig 2:**
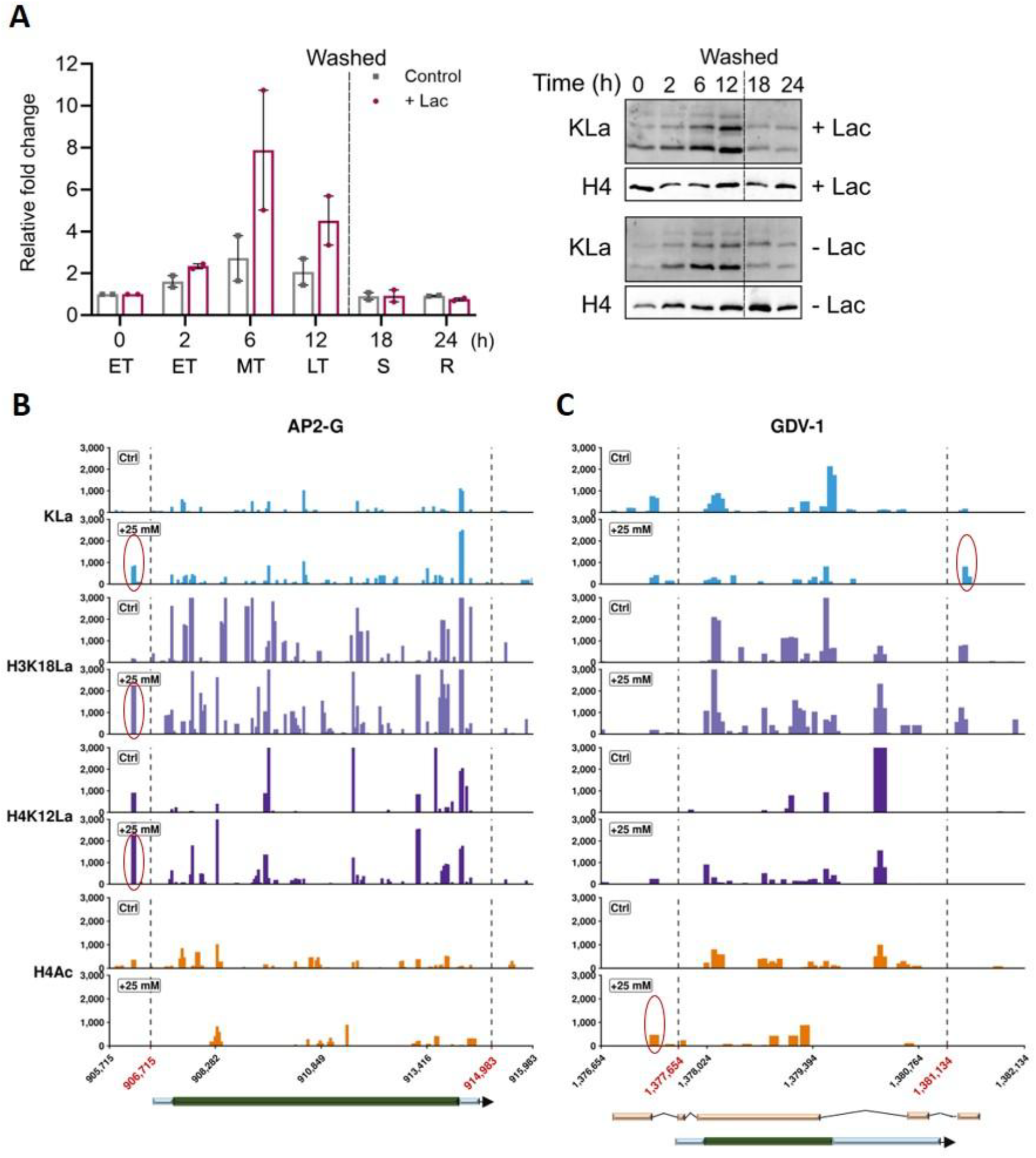
L-lactate exposure swiftly increases histone lactylation upstream of the *AP2-G* gene. A) Lysine lactylation (KLa) was measured by western blotting in parasites grown either with 25mM added L-lactate (+Lac), or in control media (Ctrl), for 12h. Both cultures were then washed into fresh media and monitored for a further 12h. Samples were taken for western blotting at 2, 6, 12, 18 and 24h. Total histone H4 was blotted at each timepoint as a control. Western blots from 2 biological replicates of this timecourse are shown; the graph shows quantification of each KLa signal relative to histone H4 in both timecourses. Stage of the parasite culture is indicated on the x-axis: ET, early trophozoite; MT, mid trophozoite; LT, late trophozoite; S, schizont; R, ring. B, C) CUT&Tag profiles of 4 histone modifications are shown across the *AP2-G* gene PF3D7_1222600 (B) and the *GDV-1* gene PF3D7_0935400 (C). Modifications are pan-lysine lactylation (KLa), H3K18La, H4K12La and H4 acetylation (H4Ac). Profiles are from NF54 trophozoites cultured in parallel under control conditions (Ctrl) or with 12h exposure to 25mM L-lactate (+25 mM). BigWig signal tracks represent the average of three biological replicates for each histone modification and condition. Peaks strongly induced in ‘+25mM’ conditions are circled in red. Gene models are shown beneath each locus; the *GDV-1* antisense RNA is shown above the *GDV-1* gene in (C).

## Discussion

Gametocytogenesis is a major virulence phenotype in *P. falciparum*. It is regulated by environmental factors in the human host, including serum metabolites [6]. Here, we show that gametocytogenesis is regulated by L-lactate: a metabolite that fluctuates greatly in malaria patients’ blood.

Gametocytogenesis was enhanced by continuous exposure of cultured parasites to 5mM added L-lactate. Prior work had reported similar results using ∼3.1mM L-lactate plus ∼5.1mM D-lactate [8], whereas here we used L-lactate alone. There was a trend towards ‘more is better’ for gametocytogenesis between 5 and 10mM L-lactate, but asexual parasite growth was inhibited when 10mM L-lactate was maintained over more than one 48h cycle, whereas 5mM could be tolerated. Notably, in any protocol for *in vitro* gametocytogenesis, cells are exposed over several cycles to endogenously-generated L-lactate as well [8], because asexual cells are metabolically active in the culture for at least the first week, and hence a truly steady level of lactate cannot be maintained. It was likewise difficult to separate enhanced maturation from enhanced commitment, but commitment was *not* enhanced by simply exposing a low-parasitaemia culture to 2 cycles of growth in 5mM added L-lactate, so lactate probably synergises with other metabolites produced during culture overgrowth.

*In vivo*, asymptomatic malaria patients without overt hyperlactataemia can still generate gametocytes [18]. Circulating blood normally contains ∼1mM lactate [19] whereas bone marrow has been reported at ∼2-3mM [8], implying that bone-marrow-sequestered parasites might experience steadily elevated lactate. However, more recent studies have argued that this is restricted to leukemic conditions [20, 21]. Circulating parasites must also experience lactate fluctuations in hypoxic organs like active muscles and in organs where sequestration is impeding blood circulation. The clear difficulty of studying this in a whole-human context makes *in vitro* modelling valuable, albeit imperfect.

*In vitro*, we clearly linked L-lactate exposure to lactylation of parasite histones and increased expression of gametocytogenesis genes. At the chromatin level, histones were inducibly lactylated upstream of the *AP2-G* gene, which is also linked to methyl-dependent epigenetic silencing via S-adenosylmethionine [6]. Indeed, the key region for methylation [22] overlapped with the lactylated region detected here. Hence, the histone code controlling *AP2-G* probably involves both methylation and lactylation (Fig 3). Control of *GDV-1*, which lies upstream of *AP2-*G, may differ: its repression depends on antisense RNA, and could be quite independent of histone marks, but intriguingly we did detect a minor lactyl peak at the end of the antisense RNA as well. Overall, this report adds lactylation to methylation in the epigenetic landscape that regulates gametocytogenesis.

**Fig 3:**
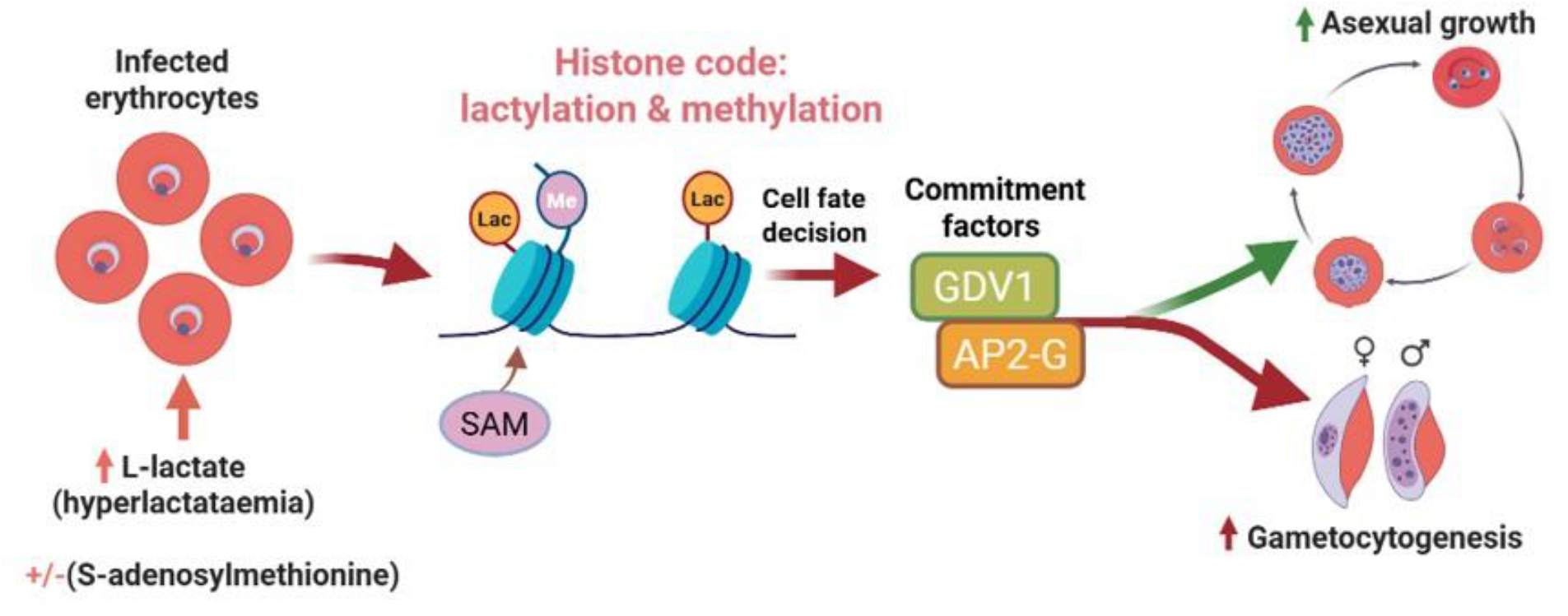
Model showing the histone code of methylation and lactylation that may control key genes that regulate gametocytogenesis.

## Methods

### Parasite culture and gametocyte conversion

The *P. falciparum* NF54 strain was maintained and synchronised to a 2h window as previously described [10]. Synchronous ring-stage cultures were seeded at 0.5% or 4% parasitaemia in 6-well plates at 4% haematocrit using a total volume of 5ml complete medium. L-lactate was then added at 0, 5 or 10 mM, and media replaced every 24h, supplemented as above, until day-16. Thin blood smears were used to count mature stage V gametocytes on days 15 and 16. For the pre-treatment experiment, synchronous ring-stage parasites were seeded at 0.5% parasitaemia and exposed to 0, 5 or 10mM L-lactate for two asexual cycles (splitting after cycle-1) before induction +/-5mM L-lactate as above. At 32 and 44hpi after 0 or 5mM lactate treatment, samples were collected for RNA extraction.

### Gene expression analysis

RNA was extracted from parasites using TRIzol Reagent (Invitrogen), purified using the Quick-RNA Miniprep kit with on-column DNase I treatment (Zymo Research), then reverse-transcribed using the SensiFAST cDNA Synthesis Kit (Meridian Bioscience). cDNA was checked for gDNA contamination as in [23], then subjected to RT-qPCR analysis using primers and cycling conditions described in Table S1.

### Western blotting

Lactate-treatment timecourses and resultant western blots were generated and analysed as in [10].

### CUT&Tag data analysis

CUT&Tag data were generated and analysed genome-wide in [17]: data from selected genes are extracted and plotted here. Normalized BigWig coverage tracks, averaged across three biological replicates for each histone mark and condition, were used to plot the genomic regions encompassing *AP2-G* (PF3D7_1222600) and *GDV-1 (*PF3D7_0935400). For visualization, BigWig signal was extracted across each locus and summarized in 25-bp genomic bins.

## Acknowledgements

This work was funded by Wellcome Discovery Award 225171/Z/22/Z to CJM. The funders had no role in study design, data collection, interpretation or the decision to submit the work for publication.

## Supporting information

**Table S1.**
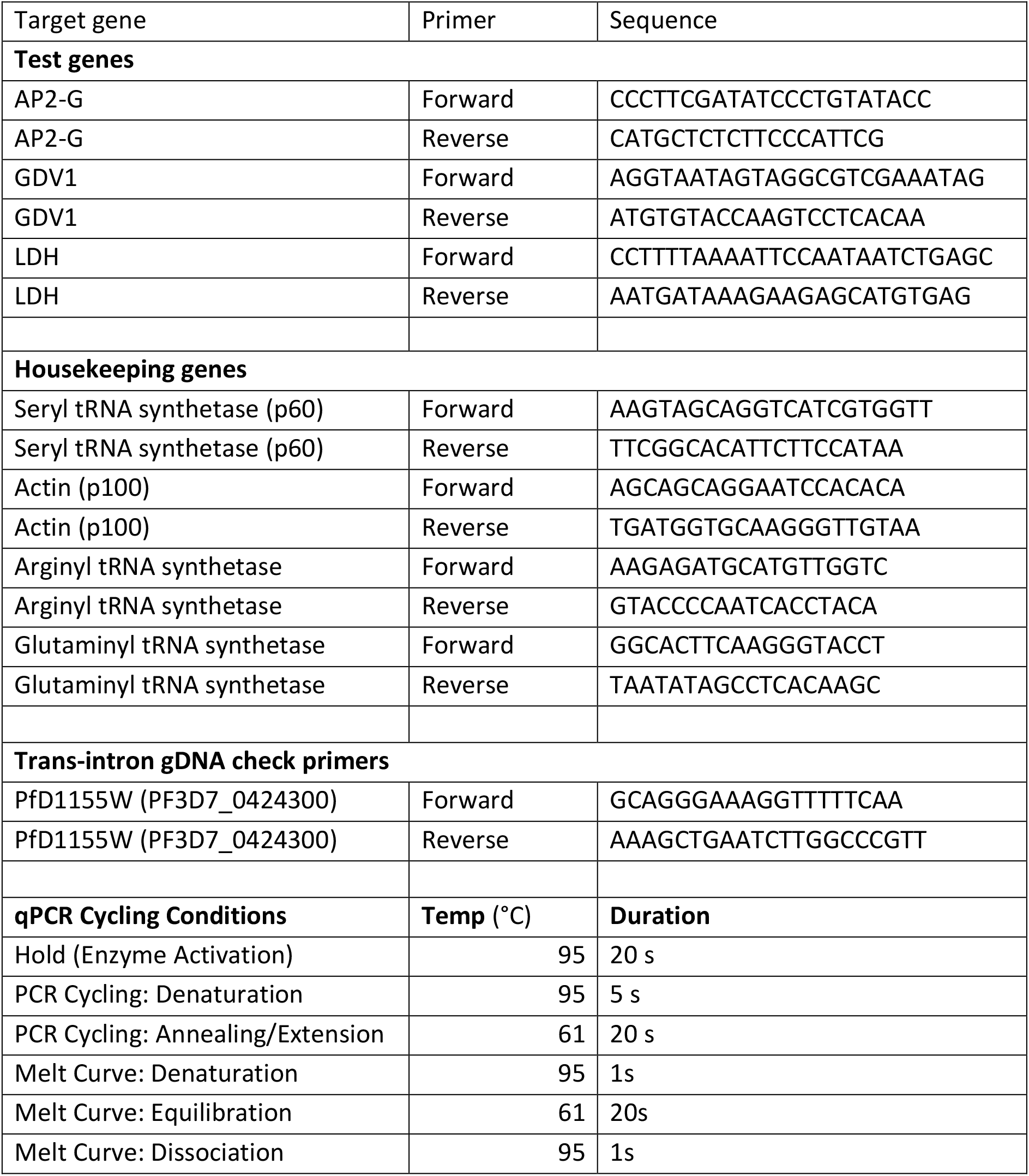
Primers and PCR conditions used in RT-qPCR.

## References

1. WHO. World Malaria Report 2023. 2023. Epub 2023.

2. Joice R, Nilsson SK, Montgomery J, Dankwa S, Egan E, Morahan B, et al. Plasmodium falciparum transmission stages accumulate in the human bone marrow. Sci Transl Med. 2014;6(244):244re5. doi: 10.1126/scitranslmed.3008882. PubMed PMID: 25009232; PubMed Central PMCID: PMCPMC4175394.

3. Aguilar R, Magallon-Tejada A, Achtman AH, Moraleda C, Joice R, Cistero P, et al. Molecular evidence for the localization of Plasmodium falciparum immature gametocytes in bone marrow. Blood. 2014;123(7):959–66. doi: 10.1182/blood-2013-08-520767. PubMed PMID: 24335496; PubMed Central PMCID: PMCPMC4067503.

4. Kafsack BF, Rovira-Graells N, Clark TG, Bancells C, Crowley VM, Campino SG, et al. A transcriptional switch underlies commitment to sexual development in malaria parasites. Nature. 2014;507(7491):248–52. doi: 10.1038/nature12920. PubMed PMID: 24572369; PubMed Central PMCID: PMCPMC4040541.

5. Sinha A, Hughes KR, Modrzynska KK, Otto TD, Pfander C, Dickens NJ, et al. A cascade of DNA-binding proteins for sexual commitment and development in Plasmodium. Nature. 2014;507(7491):253–7. doi: 10.1038/nature12970. PubMed PMID: 24572359; PubMed Central PMCID: PMCPMC4105895.

6. Harris CT, Tong X, Campelo R, Marreiros IM, Vanheer LN, Nahiyaan N, et al. Sexual differentiation in human malaria parasites is regulated by competition between phospholipid metabolism and histone methylation. Nat Microbiol. 2023;8(7):1280–92. doi: 10.1038/s41564-023-01396-w. PubMed PMID: 37277533.

7. Neveu G, Beri D, Kafsack BF. Metabolic regulation of sexual commitment in Plasmodium falciparum. Curr Opin Microbiol. 2020;58:93–8. doi: 10.1016/j.mib.2020.09.004. PubMed PMID: 33053503; PubMed Central PMCID: PMCPMC7746583.

8. West R, Sullivan DJ. Lactic acid supplementation increases quantity and quality of gametocytes in Plasmodium falciparum culture. Infection and immunity. 2020;15(89(1)):e00635–20. doi: 10.1128/IAI.00635-20. PubMed PMID: 33077626.

9. Merrick CJ. Histone lactylation: a new epigenetic axis for host-parasite signalling in malaria? Trends Parasitol. 2023;39(1):12–6. doi: 10.1016/j.pt.2022.10.004. PubMed PMID: 36357308.

10. Jabre I, Andoh NE, Naldoni J, Gregory W, Yoon CE, Cunnington AJ, et al. Histone lactylation: a new epigenetic mark in the malaria parasite Plasmodium. PLoS genetics. 2025;21(12):e1011991. Epub Dec 19. doi: 10.1371/journal.pgen.1011991.

11. Vavricka J, Broz P, Follprecht D, Novak J, Krouzecky A. Modern Perspective of Lactate Metabolism. Physiol Res. 2024;73(4):499–514. doi: 10.33549/physiolres.935331. PubMed PMID: 39264074; PubMed Central PMCID: PMCPMC11414593.

12. Vander Jagt DL, Hunsaker LA, Campos NM, Baack BR. D-lactate production in erythrocytes infected with Plasmodium falciparum. Molecular and biochemical parasitology. 1990;42(2):277–84. doi: 10.1016/0166-6851(90)90171-h. PubMed PMID: 2270109.

13. Merrick CJ, Huttenhower C, Buckee C, Amambua-Ngwa A, Gomez-Escobar N, Walther M, et al. Epigenetic dysregulation of virulence gene expression in severe Plasmodium falciparum malaria. The Journal of infectious diseases. 2012;205(10):1593–600. doi: 10.1093/infdis/jis239. PubMed PMID: 22448008; PubMed Central PMCID: PMCPMC3415821.

14. Filarsky M, Fraschka SA, Niederwieser I, Brancucci NMB, Carrington E, Carrio E, et al. GDV1 induces sexual commitment of malaria parasites by antagonizing HP1-dependent gene silencing. Science 2018;359(6381):1259–63. doi: 10.1126/science.aan6042. PubMed PMID: 29590075; PubMed Central PMCID: PMCPMC6219702.

15. Prajapati SK, Dong JX, Morahan BJ, Dotrang T, Barbeau MC, Williams AE, et al. A single valine to leucine switch disrupts Plasmodium falciparum AP2-G DNA binding and reveals GDV1’s role in ap2-g activation. Nature communications. 2026;17(1):1719. doi: 10.1038/s41467-026-68416-1. PubMed PMID: 41547993; PubMed Central PMCID: PMCPMC12913604.

16. Zhang D, Tang Z, Huang H, Zhou G, Cui C, Weng Y, et al. Metabolic regulation of gene expression by histone lactylation. Nature. 2019;574(7779):575–80. doi: 10.1038/s41586-019-1678-1. PubMed PMID: 31645732; PubMed Central PMCID: PMCPMC6818755.

17. Jabre I, Andoh NE, Mbye H, Ngwa-Amambua A, Merrick CJ. Lactylated histones mark virulence gene families in the malaria parasite Plasmodium falciparum. BioRxiv. 2026. Epub 13 August 2026. doi.org/10.64898/2026.08.12.744368.

18. Soumare HM, Blanken SL, Ahmad A, Ooko M, Gaye PM, Jadama L, et al. Asymptomatic school children and adults are important for the human infectious reservoir for Plasmodium falciparum malaria in an area of low endemicity in The Gambia. J Infect. 2025;91(1):106507. doi: 10.1016/j.jinf.2025.106507. PubMed PMID: 40404104; PubMed Central PMCID: PMCPMC12170349.

19. Brooks GA. The Science and Translation of Lactate Shuttle Theory. Cell Metab. 2018;27(4):757–85. doi: 10.1016/j.cmet.2018.03.008. PubMed PMID: 29617642.

20. Soto CA, Lesch ML, Munger JC, Frisch BJ. Effects of Elevated Lactate in the Bone Marrow Microenvironment during Acute Myeloid Leukemia. Blood. 2022;140(Supplement 1):8620–1. doi: 10.1182/blood-2022-170125.

21. Soto CA, Lesch ML, Becker JL, Sharipol A, Khan A, Schafer XL, et al. Elevated Lactate in the AML Bone Marrow Microenvironment Polarizes Leukemia-Associated Macrophages via GPR81 Signaling. BioRxiv. 2025. doi: 10.1101/2023.11.13.566874.

22. Nakashima M, Iwanaga S, Mori T. Identification of a sequence element regulating H3K9 methylation at the ap2-g locus in Plasmodium falciparum. Sci Rep. 2026. doi: 10.1038/s41598-026-59569-6. PubMed PMID: 42443261.

23. Merrick CJ, Dzikowski R, Imamura H, Chuang J, Deitsch K, Duraisingh MT. The effect of Plasmodium falciparum Sir2a histone deacetylase on clonal and longitudinal variation in expression of the var family of virulence genes. International journal for parasitology. 2010;40(1):35–43. PubMed PMID: 19666023.

